# Administration of fusion cytokines induces tumor regression and systemic antitumor immunity

**DOI:** 10.1101/2020.02.09.940379

**Authors:** Jinyu Zhang, Xuan Zhao

**Author notes:** To whom correspondence should be addressed: Jinyu Zhang, Mianyi Biotech Corporation, Xiyong Road, Shapingba district, Chongqing, 401332, China. **One Sentence Summary:** Fusion cytokines can induce tumor regression in mice.

## Abstract

The curative effects of cancer immunotherapy are hard to be improved in solid tumors. Cytokines, as powerful immune regulators, show potential in awaking host antitumor immunity. We have previously found that administration of certain cytokine combinations induced complete tumor clearance. Here we constructed the cognate fusion cytokines and evaluated their antitumor effects in various mouse tumor models. *In situ* induced expression of the fusion cytokine IL12IL2GMCSF led to tumor eradication, even those in high advanced stage. An immune memory against other irrelated syngeneic tumors was elicited. Flow cytometry analysis revealed that tumor infiltrating CD3+ cells greatly increased, accompanied with an elevation of CD8+/CD4+ ratio. The fusion protein exhibited superior immune activating capability to cytokine mixtures in vitro, and induced tumor regression in various immune competent tumor models by intratumoral injection. To improve translational potential, an immunocytokine IL12IL2DiaNFGMCSF for systemic administration was constructed by inserting tumor targeting diabody. The protein also displayed good activities in vitro. Intravenous infusion of IL12IL2DiaNFGMCSF induced a tumor infiltrating immune cell alteration like IL12IL2GMCSF, with moderate serum IFNγ increment. Therapeutic effects were observed in various tumor models after systemic administration of IL12IL2DiaNFGMCSF, with slight toxicity. These results provide the feasibility of developing a versatile cancer immunotherapy remedy.

## Introduction

In 2018, near 10 million people died from malignant tumors worldwide (1). Cancer immunotherapy has brought new hopes for treating these fatal diseases (2). Among the various emerging therapeutics, CAR-T and immune checkpoint are two most promising remedies (3, 4). They respectively exhibited good effects in hematologic malignancies and solid tumors (5–8). However, large efforts brought little advance in further improving the therapeutic outcomes. Large amount of clinical evidences suggests that only a small partial of patients can benefit from immune checkpoint inhibition (9, 10), in which the antitumor response is long lasting. The situation is different in CART therapy. T cell infusion leads to disease remission in most of the patients, but the refractory rate is also high (11, 12). To solve these problems, immunotherapy, in combination with other modalities, is tested in various cancers (13–16). Although combination therapies improve the outcomes in some experiments, many clinical trials failed due to unreasonable design derived from incomplete understanding of the rationale of tumor immunity (17–19).

Cytokines are important molecules that regulate the immune network of the body (20, 21). They are very powerful, due to low concentration of cytokines is enough to induce a strong immune reaction. Upon stimulation, an immune cell secrets some cytokines to influence other cells. Meanwhile, it is affected by the cytokines from other cells. Like language in animals, cytokines play a role in the crosstalk of immune cells, orchestrating a specific immune response (22–24). However, their work mode is similar to neutral network, making it hard to be understood and controlled. A cytokine may display controversy functions in different immune context. It is necessary to consider the indications and microenvironment before the powerful weapon is attempted in disease treatment.

In cancer therapy, some cytokines, alone or in combinations, have exhibited good therapeutic effects in experimental animals (25–27). However, the clinical outcomes are not satisfactory in toxicity windows. Though it looks reachable to harness cytokines to defeat tumors, the severe side effects restrict the clinical application (28, 29). Some cytokines, e.g. IL12 and IL2, have great potential in enhancing cytotoxic cellular immunity (30), which is considered necessary for tumor eradication (31, 32). Because malignant tumors confront the immune system by a variety of tricks, the strategies of treating cancer on the avail of these mighty molecules are worth of deep exploration.

In a previous study, we found that some combinations of triple cytokines exhibited unexpected antitumor activities after local administration at tumor site, which is mediated by the activation of the immune system (33). As a mixture of biomacromolecules, it is hard to be translated into standard pharmaceutical manufacturing and clinical usage. Here we designed the fusion proteins of these cytokines and evaluated their therapeutic effects in mouse tumor models. It demonstrates the potential of translating the preclinical modality into human malignant tumor therapy.

## Results

### Induced expression of fusion cytokine dcIL12IL2GMCSF led to tumor regression and antitumor immunity

Firstly, we constructed an inducible B16F10 cell line to evaluate the antitumor capability of the fusion protein, dcIL12IL2GMCSF, which was a heterodimer comprising of two polypeptides ligated with disulfide bonds between IL12p35 and IL12p40 subunits (Fig. 1A). The expression of the protein could be effectively induced by dox addition, with a low background level (Fig. 1B). The tumor cells were subcutaneously inoculated into the flank of mice and induction was initiated as tumors reached indicated sizes. The expression of dcIL12IL2GMCSF completely erased the tumors less than 15mm in diameter. For large tumors with a diameter over 15mm, 40% mice died within 3 days post dox addition. However, the remaining mice successfully eliminated the large tumors and gained long term survival (Fig. 1C). There were vitiligo symptoms at original tumor sites in cured mice (Fig. 1D). Notably, no systemic vitiligo was observed even 12 months after tumor regression. The cured mice were respectively rechallenged with parental B16F10 cells, or irrelevant syngeneic tumor cells, LLC and EL4 cells, over 10 months post initial treatment. Except for one mouse in LLC group, all other mice absolutely rejected the inoculated malignant cells (Fig. 1E). It indicated the existence of an immune memory against various tumor antigens, which derived from primary inducible B16F10 tumor eradication.

**Fig. 1.**
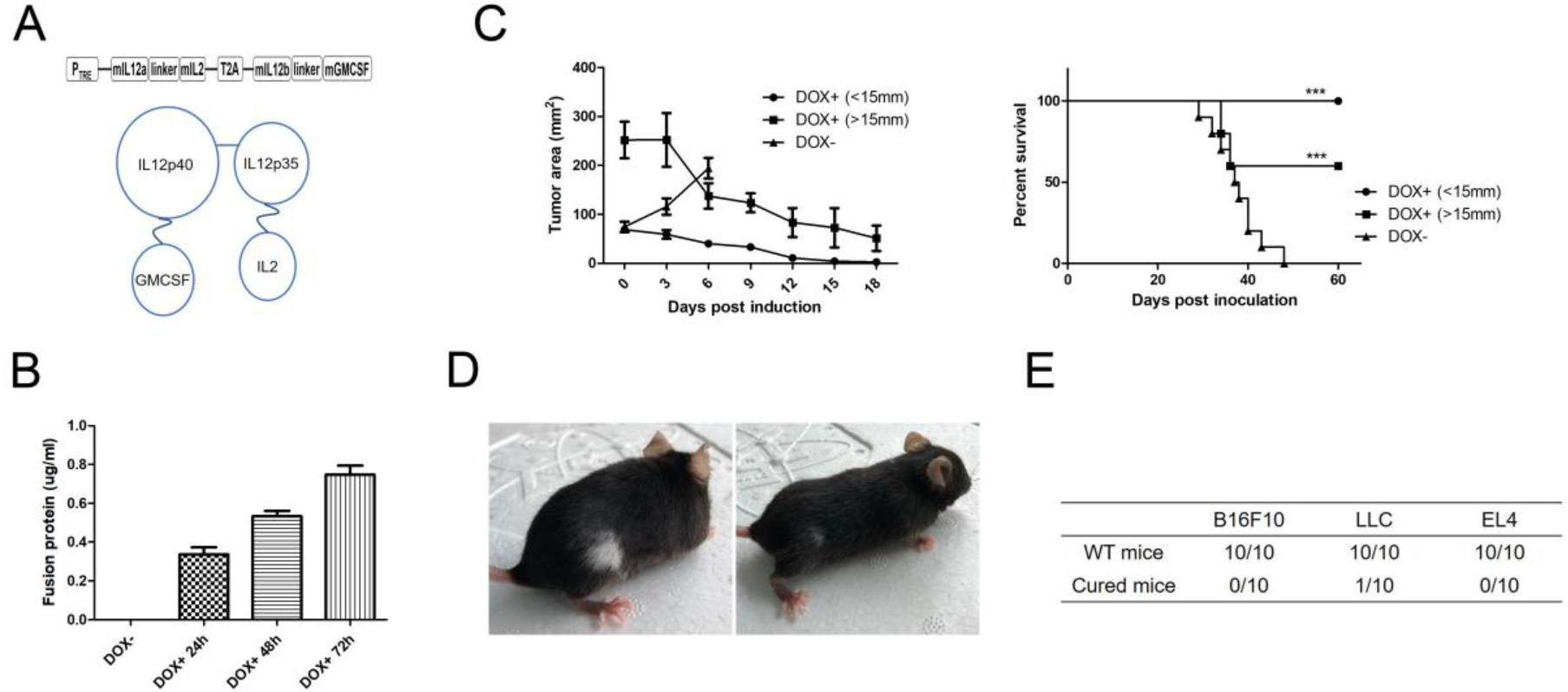
Induced expression of dcIL12IL2GMCSF led to tumor regression and antitumor immune memory. (A) Schematic representation of dcIL12IL2GMCSF expression cassette and molecule conformation. (B) The expression of dcIL12IL2GMCSF was induced by dox addition. At different time points, the supernatants were collected and subjected to ELISA measurement of fusion cytokine level. n=3. (C) The inducible B16F10 cells were subcutaneously inoculated into the flank of C57BL/6 mice. Dox was administered via drinking water when tumors reached indicated sizes. Tumor growth (n=5) and overall survival (n=10) were recorded. ***p<0.001. The experiments were repeated three times. (D) Representative photos of vitiligo occurred in cured mice. (E) The cured mice were subcutaneously injected with B16F10, LLC or EL4 tumor cells 10 months post original tumor inoculation. Age matched wild type mice were used as control. Numbers in the table indicated death/inoculated mice.

### Immune cell alterations induced by dcIL12IL2GMCSF expression

Considering the roles of IL12, IL2 and GMCSF in the immune system, we explored the alteration of immune cells during the antitumor response. Splenocytes and tumor infiltrating lymphocytes were collected and subjected to FACS analysis at different times post induction. In spleen, NK1.1+ and CD3+ subset cells gradually decreased (Fig. 2A). But the CD8+/CD4+ ratio of T cells was not influenced (Fig. 2B). In tumors, NK1.1+ cells reduced, but the CD3+ cells greatly increased at day 3 post induction (Fig. 2A). Especially, CD8+/CD4+ ratio of T cells clearly elevated, indicating a cytotoxic T cell infiltration in tumors (Fig. 2B). Intracellular IFNγ staining demonstrated the existence of activated CD8+ T cells (Fig. 2D). CD11c+ cells gradually decreased in tumors, and increased in spleen (Fig. 2C). The change of CD11b+ cells were similar with CD11c+ population, a decline in tumor and a rise in spleen. Moreover, CD11b+MHCII+ subset in spleen markedly raised post induction, suggesting an improved antigen presentation capability of these cells (fig. S1A). There was slight increase of activated B220+ cells in spleen, and the ratio in tumors significantly increased (fig. S1B). In addition, we found that the regulatory cell receptor PD1 was greatly unregulated in T cells (fig. S1C,D,E). At day 3 post induction, most of tumor infiltrating T cells were PD1 positive, suggesting a powerful negative immune regulation mechanism in antitumor immunity.

**Fig. 2.**
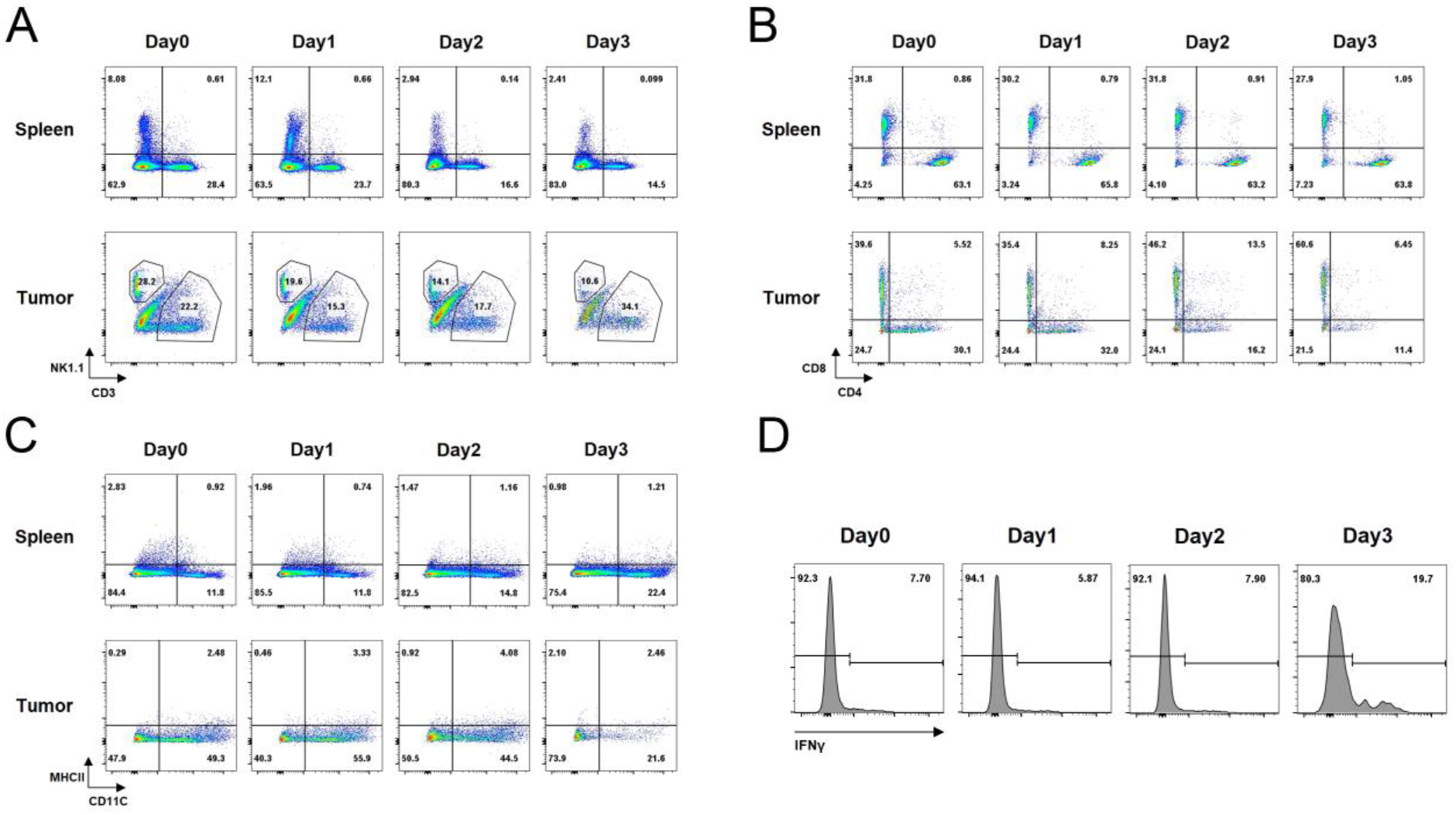
The alterations of immune cells during induced expression of dcIL12IL2GMCSF. The inducible B16 cells were subcutaneously inoculated into the flank of C57BL/6 mice. At different times post dox administration, splenocytes and tumor infiltrating lymphocytes were isolated and subjected to flow cytometry analysis. The gated CD45+ cells were analyzed with the groups of NK1.1, CD3 (A) or CD11C, MHCII (C). CD4 and CD8 expression were analyzed in gated CD3+ cells (B). Graph D showed the intracellular staining of IFNγ in CD3+/CD8+ cells from splenocytes. The experiments were performed twice with n=2 for each run.

### The fusion cytokine scIL12IL2GMCSF exhibited superior immune activation capability in vitro and antitumor effects in vivo

In previous studies, the expression of IL12 from bi-cistron construction was low. Thus we designed a single cistron cassette to improve the productivity of IL12IL2GMCSF fusion protein (Fig. 3A). ELISA measurement demonstrated that the expression of scIL12IL2GMCSF was much higher than dcIL12IL2GMCSF, reaching 100 ug/ml (fig. S2). The fusion protein could be easily purified using affinity magnetic beads (Fig. 3B). Next the activities of the fusion protein were detected by stimulating the IFNγ secretion from splenocytes. The activation capability of scIL12IL2GMCSF was superior to the mixture of equivalent IL12, IL2 and GMCSF. At low concentration, little activity of scIL12IL2GMCSF was detected. However, the fusion protein exhibited much higher activity than other groups at high concentration (Fig. 3C). Subsequently, the therapeutic potential of scIL12IL2GMCSF was evaluated by intratumoral injection.

**Fig. 3.**
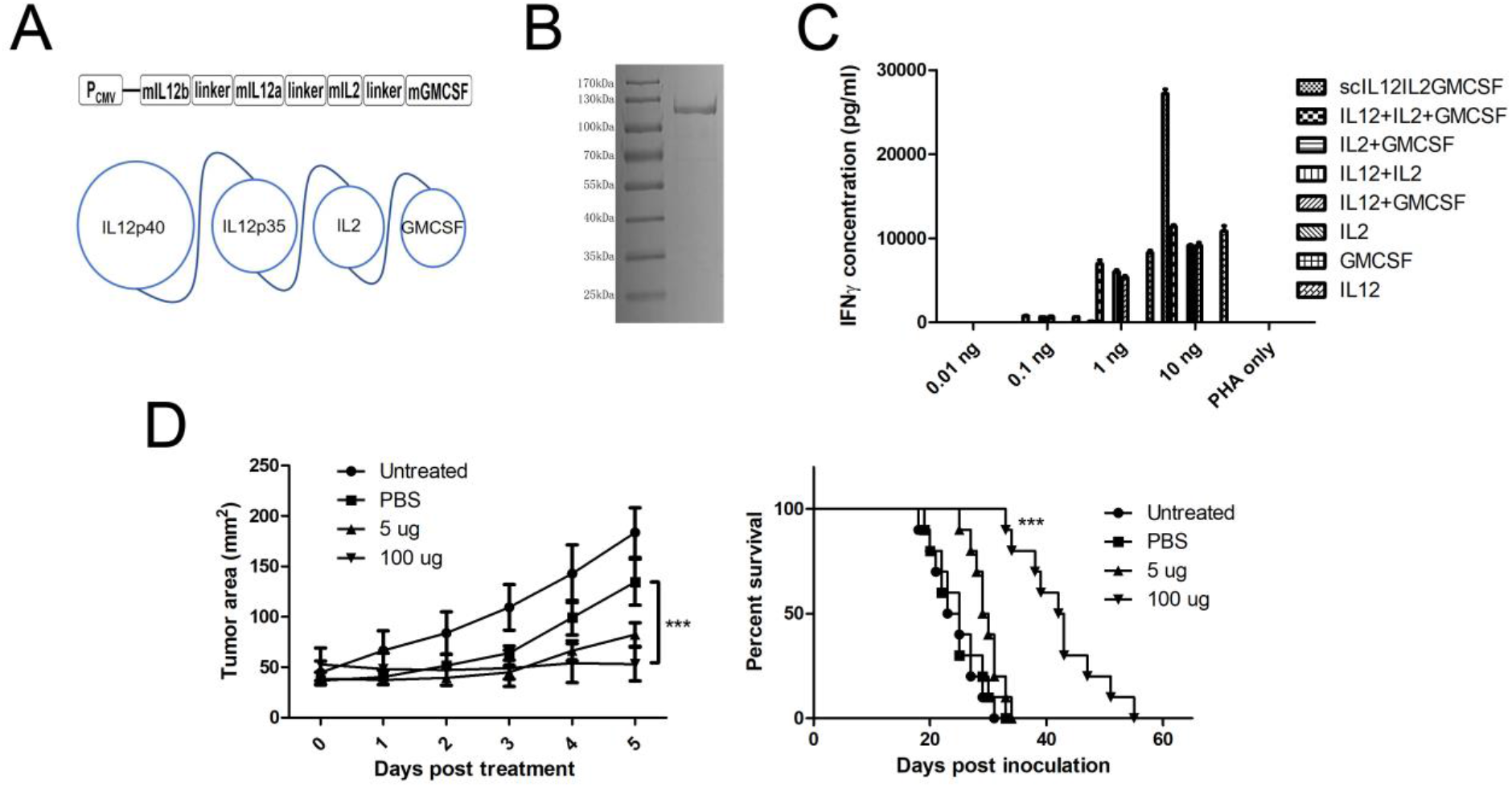
The immune activation capability and antitumor effects of fusion cytokine scIL12IL2GMCSF. (A) Schematic representation of scIL12IL2GMCSF expression cassette and molecule conformation. (B) SDS page electrophoresis of purified scIL12IL2GMCSF protein. (C) Splenocytes from C57BL/6 mice were plated into 96-well plate and stimulated with indicated cytokines plus PHA for 24 hours. IFNγ in the supernatants was measured by ELISA. n=3. (D) B16F10 cells were subcutaneously inoculated into the flank of C57BL/6 mice. scIL12IL2GMCSF or PBS was intratumorally injected into the lesions when tumors reached 5-9 mm in diameter. Tumor growth (n=5) and overall survival (n=10) were recorded. ***p<0.001. These experiments were repeated twice.

Considering the influence of solvent on treatment, we compared the effect of carboxymethyl cellulose, chitosan and glycerol. The latter two both displayed inhibition activities (fig. S3). Glycerol was selected for following usage due to it was an approved excipient in clinic. The administration of scIL12IL2GMCSF significantly suppressed subcutaneous B16F10 melanoma growth. The therapeutic outcome was dose related, and high dose protein greatly improved the survival of tumor bearing mice (Fig. 3D). Some weight loss was observed during treatment, which was drug irrelated because injection of PBS also led to this effect (fig. S4). Interestingly, administration of low dose scIL12IL2GMCSF completely eradicated subcutaneous B16F10-rtTA tumors (fig. S5). It was likely due to that rtTA protein provided a predominant neoantigen for effective immune recognition.

### Intratumoral injection of scIL12IL2GMCSF induced tumor clearance in various tumor models

Based on the antitumor activity in melanoma, we additionally tested the curative effects of scIL12IL2GMCSF in several other mouse malignant tumors. For LLC (Fig. 4A), EL4 (Fig. 4B) and CT26 (Fig. 4C) tumors, single injection induced complete tumor regression without recurrence. There was no death in scIL12IL2GMCSF treatment group. For 4T1 tumor (Fig. 4D), one mouse died from tumor relapse, though the original tumor size reduced after drug injection. In general, intratumoral injection of scIL12IL2GMCSF exhibited curative potential in tested mouse tumor models.

**Fig. 4.**
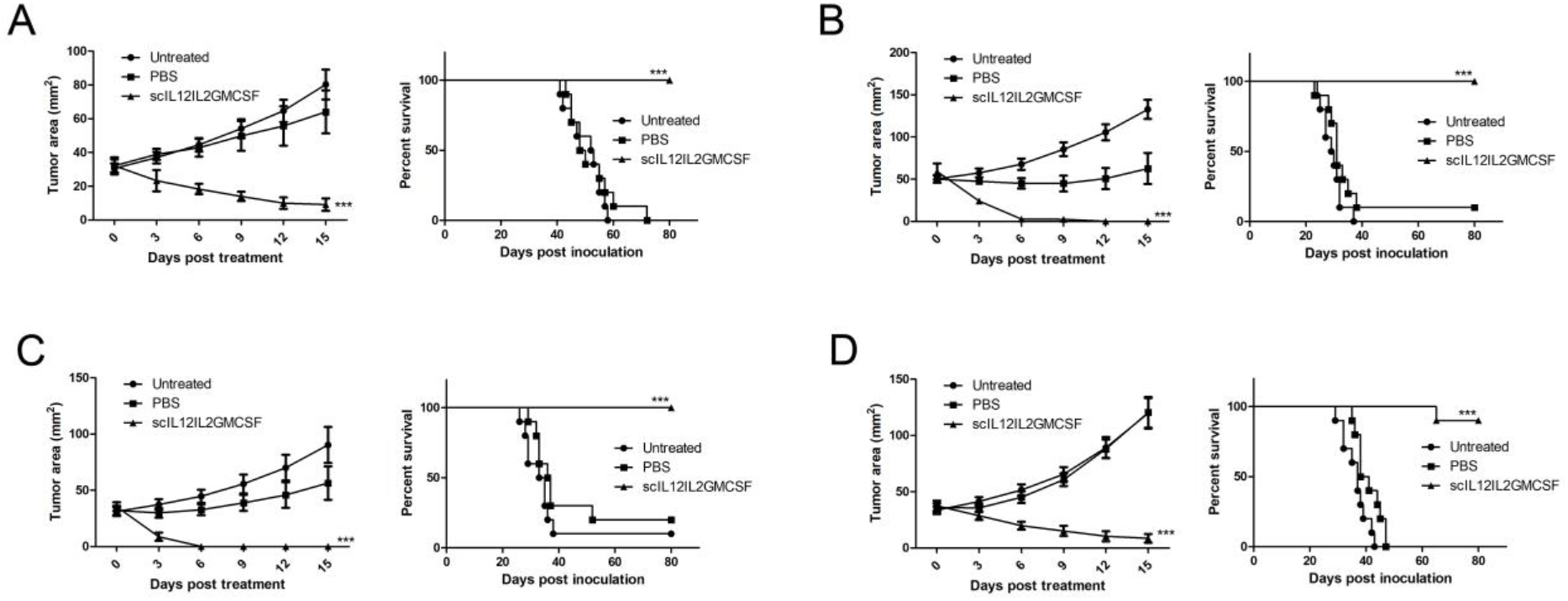
Therapeutic effects of intratumoral administration of scIL12IL2GMCSF in various mouse tumor models. Mouse LLC (A) or EL4 (B) tumor cells were subcutaneously inoculated into the flank of C57BL/6 mice. Mouse CT26 (C) or 4T1 (D) tumor cells were subcutaneously inoculated into the flank of Balb/c mice. When tumor reached 5-9 mm in diameter, 100 ug scIL12IL2GMCSF or PBS was intratumorally injected into the lesions. Tumor growth (n=5) and overall survival (n=10) were recorded. ***p<0.001. These experiments were repeated twice.

### The immunocytokine scIL12IL2DiaNFGMCSF induced IFNγ expression and immune cell alterations

To improve the translational application of this remedy, we designed the intravenous formulation of the fusion cytokine by adding a tumor targeting diabody DiaNF (Fig. 5A), which consisted of scFv from F8 and NHS76 antibodies. In vitro activity measurement revealed that the IFNγ stimulating capability of scIL12IL2DiaNFGMCSF is higher than scIL12IL2GMCSF (Fig. 5B). After intravenous injection, the tumor tissues were collected and subjected to western blot analysis. Compared with control, a band of ~75 kDa was clearly detected in tumors one day post injection, which might be IL12-IL2 fragment released from scIL12IL2DiaNFGMCSF by thrombin cleavage (Fig. 5C). Then we explored the effects of fusion cytokine administration on the host immune system. After infusion, serum IFNγ clearly elevated and rapidly declined to low level within two days, indicating a transient systemic immune activation (Fig. 5D). It also led to some immune cell alterations detected by FACS. There was a significant increase of tumor infiltrating CD3+ cells (Fig. 5E), in which CD8+/CD4+ ratio greatly raised (fig. S6). T cells in spleen were slightly decreased, without changes in CD8+/CD4+ ratio (Fig. 5E, fig. S6). CD11c+ cells increased both in tumor and spleen, and the intratumor cells seemed activated due to the enhanced expression of MHCII molecule (Fig. 5E). Although no changes observed in spleen, like CD11c+ cells, CD11b+ cells were activated in tumors with an elevated MHCII expression (fig. S6). These results suggested that intravenous administration of scIL12IL2DiaNFGMCSF efficiently induced an immune activation, especially in tumors.

**Fig. 5.**
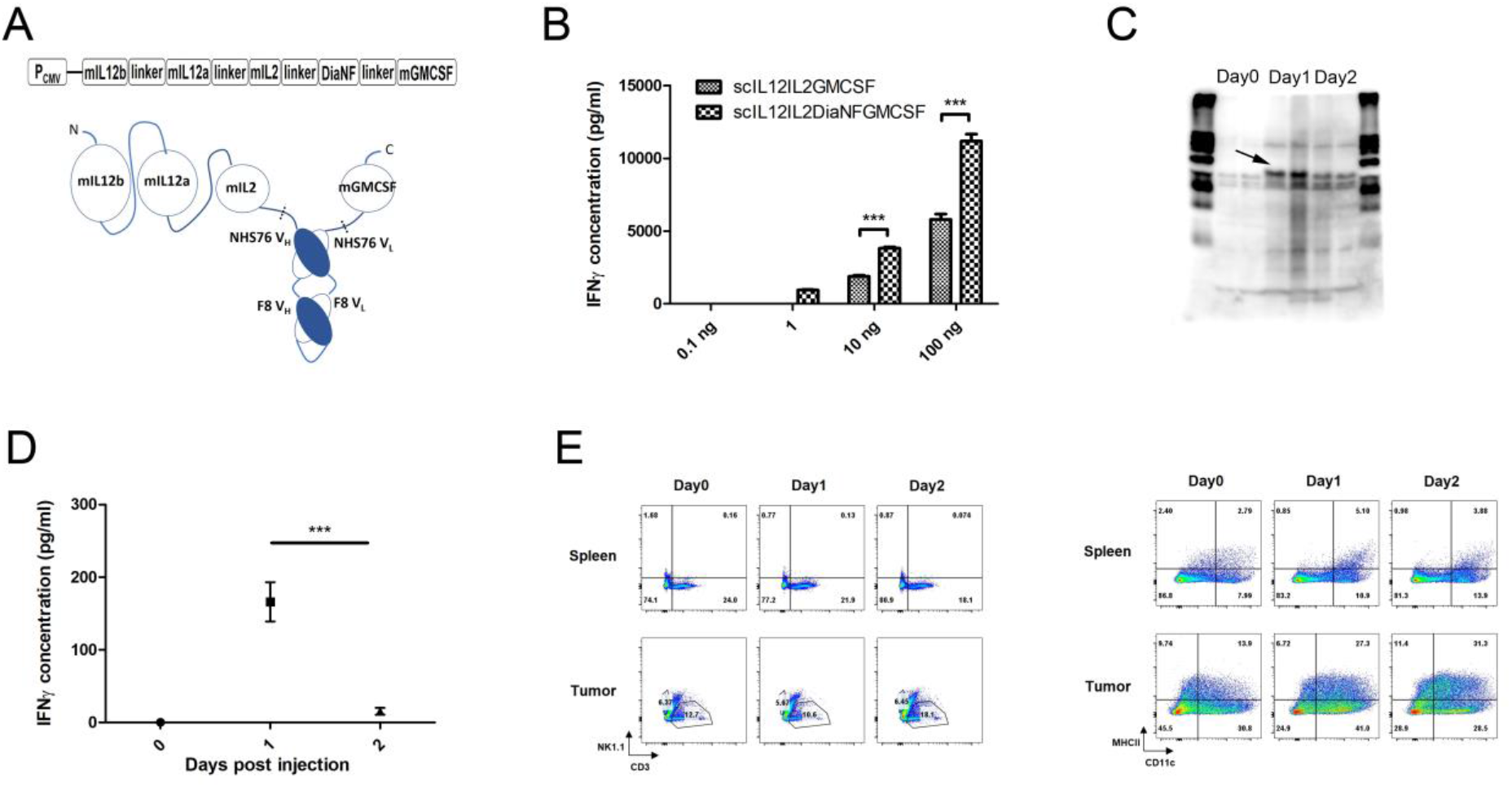
The immunocytokine scIL12IL2DiaNFGMCSF stimulated immune activation in vitro and in vivo. (A) Schematic representation of scIL12IL2DiaNFGMCSF expression cassette and molecule conformation. The dashed lines indicated thrombin cleavage sites. (B) Splenocytes from C57BL/6 mice were plated into 96-well plate and stimulated with scIL12IL2GMCSF or scIL12IL2DiaNFGMCSF plus PHA for 12 hours. IFNγ in the supernatants was measured by ELISA. n=3. (C) Mouse LLC tumor cells were subcutaneously inoculated into the flank of C57BL/6 mice. When tumor reached 5-8 mm in diameter, 200 ug scIL12IL2DiaNFGMCSF was intravenously injected into mice. Tumor tissues were separately collected at day 0, 1, or 2 post injection, and subjected to western blot analysis, using anti-IL12p40 antibody. Arrow indicated distinctive bands ~75kd. (D and E) Mouse LLC tumor cells were subcutaneously inoculated into the flank of C57BL/6 mice. When tumor reached 5-8 mm in diameter, 200 ug scIL12IL2DiaNFGMCSF was intravenously injected into mice. At day 0, 1 or 2 post injection, the serums were isolated and subjected to ELISA measurement of IFNγ (D), the splenocytes and tumor infiltrating lymphocytes were subjected to flow cytometry analysis (E). The gated CD45+ cells were analyzed with the groups of NK1.1, CD3 or CD11c, MHCII. The experiments were performed twice with n=2 for each run. ***p<0.001.

### Intravenous administration of scIL12IL2DiaNFGMCSF suppressed tumor growth in various tumor models

Next, the therapeutic effects of scIL12IL2DiaNFGMCSF were evaluated in mouse tumor models. To find out the best administration schedule, the fusion cytokine was intravenously injected into LLC tumors bearing mice by every day (qd), two days (q2d) or three days (q3d), for total 5 doses. Although there was no significant difference in survival between qd and q2d group, the former displayed better effects in tumor growth suppression (Fig. 6A). Mice bearing B16F10 or MC38 tumors were treated using qd schedule. The intravenous administration greatly suppressed tumor growth and elongated survival time (Fig. 6B, 6C). Importantly, no weight loss was observed during therapeutic process (fig. S7). It indicated that the antitumor effects of the fusion cytokine could be achieved by systemic administration.

**Fig. 6.**
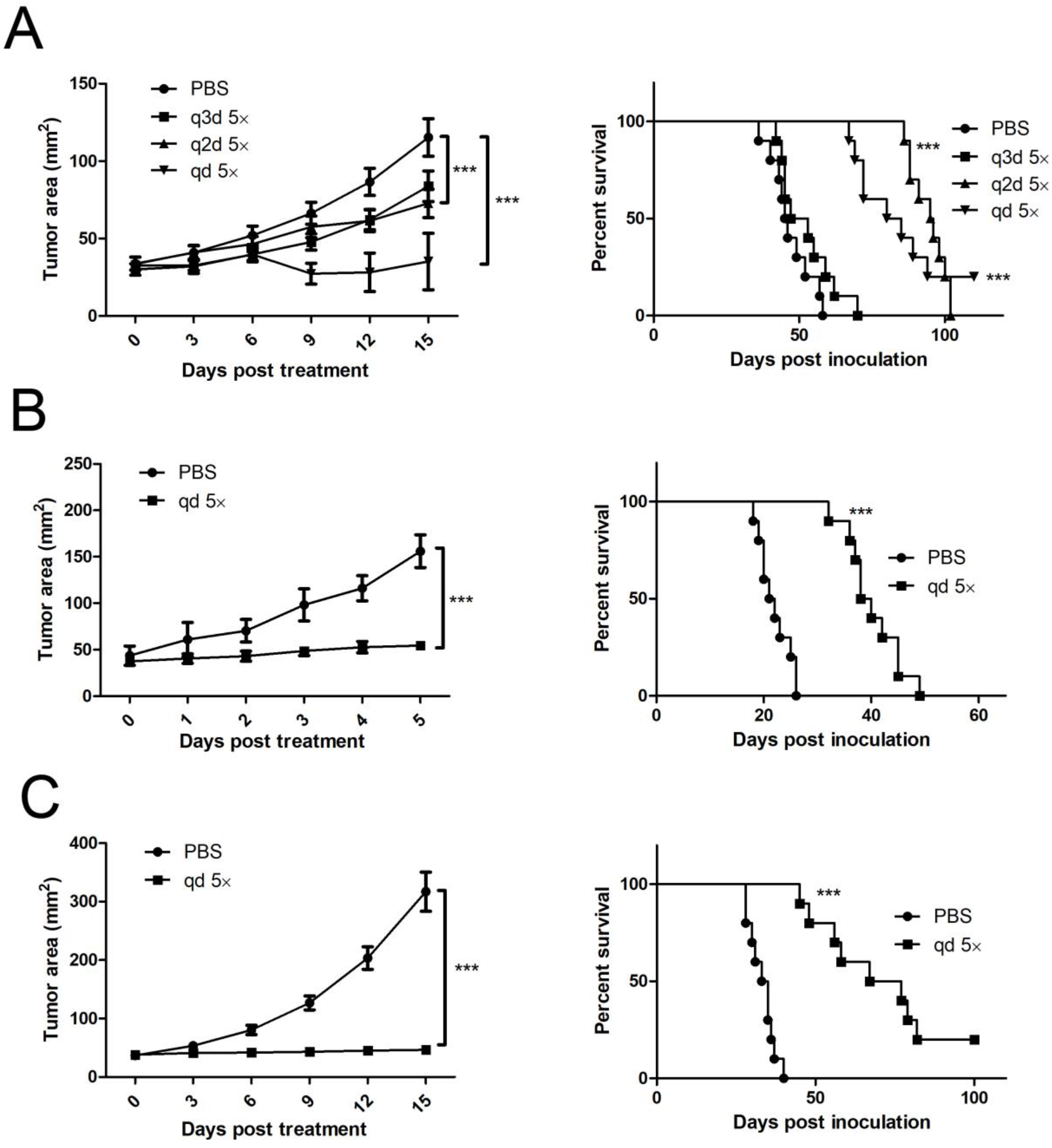
Therapeutic effects of intravenous administration of scIL12IL2DiaNFGMCSF in various mouse tumor models. (A) Mouse LLC tumor cells were subcutaneously inoculated into the flank of C57BL/6 mice. When tumor reached 5-9 mm in diameter, 50 ug scIL12IL2DiaNFGMCSF was intravenously injected every day (qd), two days (q2d) or three days (q3d), for total five doses. Tumor growth (n=5) and overall survival (n=10) were recorded. Mouse B16F10 (B) or MC38 (C) tumor cells were subcutaneously inoculated into the flank of C57BL/6 mice. When tumor reached 5-9 mm in diameter, daily intravenous administration of 50 ug scIL12IL2DiaNFGMCSF was conducted for five days. Tumor growth (n=5) and overall survival (n=10) were recorded. ***p<0.001. These experiments were repeated twice.

## Discussion

In our previous study, we have revealed some antitumor cytokine combinations, using the in vivo cytokine screen system. Among them, IL12+GMCSF+IL2 showed the most promising therapeutic potential. However, the cytokines combination is hard to be translated into pharmaceuticals, due to the expensive cost for the manufacture of three biomolecules and the difficulty in the quality control of protein mixture. Linked by peptide linkers, the fusion proteins showed comparable antitumor effects. The fusion might ameliorate the toxicity of these cytokines, because the acute death of mice bearing large tumors after induction decreased in fusion protein group. Some IL12-IL2 fusion cytokines also exhibited good safety in tumor bearing mice (34, 35). Compared to individual cytokine, one IL12IL2GMCSF fusion molecule can bind the cognate IL12, IL2 or GMCSF receptors, making it much easier to be captured and retained at tumor lesion, reducing the systemic leakage and consequent toxicity. Notably, the yield of two chain IL12IL2GMCSF fusion is only at ug/ml level, close to some other IL12 based cytokine fusions (36, 37), which is likely ascribed to homodimerization of IL12p40 subunit. The yield of single chain IL12IL2GMCSF protein reaches 100 ug/ml, providing a molecule entity much suitable to large scale manufacture.

The concentration of cytokines should be maintained at tumor site to provide a sustained immune stimulation. Many slow release materials have been attempted in local cytokine delivery (38, 39). PLGA microspheres is frequently used to encapsulate cytokines or cytokine expressing cells, and exhibit good safety and sustained release characteristics (40–42). Chitosan and alginate are also widely used in preclinical studies (43–46). However, these methods are complicated to operate or not been approved in clinic. In this work, glycerol is utilized as a carrier to deliver cytokine proteins. Firstly, the viscosity of the solution leads to the retention of drugs at injection site. Secondly, glycerol is a common preservative of purified protein, which is beneficial to maintain cytokine activities. Moreover, injection of glycerol alone exhibits certain antitumor effects, which might enhance the therapeutic effects of cytokines through a synergistic effect. Of note, administration of glycerol brings systemic toxicities, e.g. weight loss, in tumor bearing mice. It is likely due to the high dose/weight ratio in mice, and can be ameliorated in human.

Though discovered for many years, the toxicities of cytokines restrict their application in clinic. In addition to local delivery, antibody conjugated cytokine is a good choice to reduce the toxicities (47, 48). Compared to free cytokines, immunocytokines exhibit superior therapeutic effects in a variety of mouse tumors (49, 50). In a previous research, an immunocytokine carrying IL12-IL2 playload markedly suppressed the growth of Epcam-LLC tumor, indicating the potential of immunocytokine with multiple cytokine playloads (35). Alternatively spliced extra-domain A (EDA) and extra-domain B (EDB) of fibronectin, histone-DNA complex (HDC), epidermal growth factor receptor (EGFR), et al, are the frequently adopted targets in these molecule designs. Some IL-12 based immunocytokines have entered clinical studies. Among them, BC1-IL12 and NHS-IL12 are proved safe in stage I clinical research (51, 52). The maximal tolerated doses are 15 ug/kg for the former and 16.8 ug/kg for the latter. Considering the MTD of free IL12 is 0.5-1.0 ug/kg in preceding clinical studies, the fusion of antibodies to cytokines greatly ameliorates the toxicities of systemic administration. In this study, we construct a bispecific scFv, targeting histone-DNA complex and EDA. Anti-HDC is placed at inner side, because its target is free and less influenced by steric interference. Compared to single target, this design provides an additional chemotactic factor and might improve the targetability of the immunocytokine.

A great deal of clinical practice has indicated that monotherapy of cancer reaches a ceiling, including immunotherapy. Recently, some novel combinational immunotherapies exhibit excellent curative effects (53–56). These results suggest that multiple immune stimulations have the capability of eliciting immune response to erase malignant tumors. Considering the functions of IL12, IL2 and GMCSF in the immune system, it can be envisaged that a positive feedback loop plays a pivotal role in the fusion protein mediated tumor elimination (fig. S8). Briefly, IL12 and IL2 synergistically activate NK cells to kill a part of malignant cells (57, 58), especially those escape from T cell surveillance through downregulating MHC class I and β2m expression. If existence, pre-existing infiltrating T cells also participate in the antigen release process. A group of super dendritic cells with enhanced antigen presentation capability, which is acquired from the synergy of IL12 and GMCSF (59, 60), captures those tumor antigens and migrated to draining lymph nodes to prime tumor specific T cells, which include those clones against low abundant neoantigens. These T cell clones are further activated and expanded by the synergistic action of IL12 and IL2 (61, 62), and infiltrate into tumors to kill more malignant cells. The killing-presentation cycle finally generates a multiplex antitumor immune repertoire against most of heterogeneous cancer cells. Based on this mechanism, it is reasonable that mice cured from melanoma acquire resistance to other unrelated syngeneic tumors. Taken together, our research presents novel fusion proteins, which exhibit great antitumor potential and are straightforward to be translated into human cancer treatment.

## Materials and Methods

### Cell lines and mice

Mouse B16F10 melanoma, Lewis lung carcinoma (LLC), EL4 lymphoma, CT26.WT colon carcinoma, MC38 colon adenocarcinoma, human embryonic kidney cell line 293 and 293FT were cultured in Dulbecco’s modified Eagle’s Medium (DMEM) supplemented with 10% FBS (Life Technologies), penicillin/streptomycin (Life Technologies) at 37°C in 5% CO2 incubator. Mouse 4T1 mammary carcinoma cell line was cultured in RPMI1640 medium supplemented with 10% FBS (Life Technologies), penicillin/streptomycin (Life Technologies) at 37°C with 5% CO2.

C57BL/6 and BALB/c mice were purchased from Beijing Huafukang bioscience company. 8-16 weeks old mice were used for tumor experiments. All animals were raised under normal environment conditions. All animal experiments were conducted in accordance with guideline for the care and use of laboratory animals.

### Plasmid construction and cell transduction

The construction of inducible expression system was described in a previous study (33). dcIL12IL2GMCSF is a fusion protein consisting of two cistrons, IL12b-GMCSF and IL12a-IL2, which were ligated by a T2A peptide. The DNA encoding dcIL12IL2GMCSF was synthesized and subcloned into pLentis-TRE-MCS-PGK-PURO vector between BamHI and XhoI sites, generating inducible vector pLentis-TRE-dcIL12IL2GMCSF-PGK-PURO. scIL12IL2GMCSF is a fusion protein in which IL12b, IL12a, IL2 and GMCSF were tandemly ligated by (G_4_S)_3_ linkers. scIL12IL2DiaNFGMCSF is designed by inserting a heterogenous tumor targeting diabody DiaNF between IL2 and GMCSF of scIL12IL2GMCSF. Thrombin cleavage sequence LVPRGS is inserted into the linkers between DiaNF and cytokines. The DNA encoding dcIL12IL2GMCSF, scIL12IL2GMCSF and scIL12IL2DiaGMCSF were synthesized and subcloned into vector pLentis-CMV-MCS-IRES-PURO between BamHI and XhoI sites, generating expression vectors pLentis-CMV-dcIL12IL2GMCSF-IRES-PURO, pLentis-CMV-scIL12IL2GMCSF-IRES-PURO and pLentis-CMV-scIL12IL2DiaNFGMCSF-IRES-PURO. 6*His was added to the C terminal of fusion proteins to provide an affinity purification tag.

The lentiviral particles of these vectors were produced by cotransfection of pMD2.G, psPAX2 and lentiviral vectors into 293FT cells. B16F10-rtTA cells were transduced with pLentis-TRE-dcIL12IL2GMCSF-PGK-PURO virus and selected with 3 ug/ml puromycin plus 8 ug/ml blasticidin, generating doxycycline (dox) inducible cells, B16F10-rtTA(TRE-dcIL12IL2GMCSF). For protein expression, 293 cells were transduced with pLentis-CMV-dcIL12IL2GMCSF-IRES-PURO, pLentis-CMV-scIL12IL2GMCSF-IRES-PURO and pLentis-CMV-scIL12IL2DiaNFGMCSF-IRES-PURO virus and selected using 3 ug/ml puromycin, generating stable transduced cells, 293(dcIL12IL2GMCSF), 293(scIL12IL2GMCSF) and 293(scIL12IL2DiaNFGMCSF).

### Induced expression of dcIL12IL2GMCSF

B16F10-rtTA(TRE-dcIL12IL2GMCSF) cells were plated into a 24 well plate at 5×10^4^ cells/well in 700ul medium. 100 ng/ml dox was separately administered at 24, 48 or 72 hours, all supernatants were collected at 96 hours post cell plating. The concentration of fusion protein in the supernatants was measured with mouse IL12p70 ELISA Kits (Neobioscience), according to the manufacturer’s instructions.

10^5^ B16F10-rtTA(TRE-dcIL12IL2GMCSF) cells were subcutaneously injected into the right flank of mice. When tumors reached indicated size, dox was administered by adding 2 g/L dox into drinking water. The perpendicular diameters of tumors were measured using a caliper, and tumor area was calculated by long diameter×short diameter. To explore the antitumor immune memory, 10^5^ B16F10, 2×10^5^ LLC or 10^6^ EL4 tumor cells were subcutaneously injected into the left flank of cured mice, ten months post primary tumor eradication.

### Flow cytometric analysis of immune cells

Mouse spleen is minced by a syringe plunger on a 70um cell strainer. After lysis of red blood cells, the dissociated splenocytes were rinsed with PBS and resuspended in FACS buffer (PBS+2%FBS+5mM EDTA). Harvested solid tumor tissues were dissociated into single cells by mechanical disaggregation and then collagenase digestion (Dnase 1ug/ml, collagenase type II 1mg/ml, CaCl_2_ 5 mM in PBS) at 37°C for an hour. After passing through 70um cell strainer, tumor infiltrating lymphocytes (TIL) were isolated using 40:80% Percoll (GE healthcare). Prior to staining, the splenocytes and TILs were blocked for 30 minutes in FACS buffer containing Fc Blocker (1ug/ml). The cells were stained with these antibodies: mouse CD45 AF700, mouse CD3 BV421, mouse CD4 PE, mouse CD8 FITC, mouse NK1.1 APC, mouse PD1 PE/CY7, mouse CD11B APC, mouse CD11C PE, mouse MHCII FITC, mouse B220 BV510, mouse mIgD PE/CY7 and mouse IFNγ APC/CY7 (Biolegend). FVD506 and 7AAD were used to distinguish viable cells. Cells were detected by flow cytometry using a BD LSRFortessa Flow cytometer. Acquired data were analyzed using Flowjo software.

### Production of recombinant proteins

293(dcIL12IL2GMCSF), 293(scIL12IL2GMCSF) and 293(scIL12IL2DiaNFGMCSF) cells were separately inoculated into 15 cm dishes in DMEM medium plus 10% FBS. After the confluence reached 90%, culture medium was replaced with 35 ml CDM4HEK293 medium (Hyclone) supplemented with 4 mM L-Glutamine. Five days later, the supernatants were collected and filtrated through 0.45 um filter (Millipore). After concentrated to 1 ml using Amicon Ultra-15 centrifugal filter unit (Millipore), the cytokine solutions were purified using BeaverBeads IDA-Nickel Kit (Beaverbio), according to the manufacturer’s instructions. The cytokine concentration was determined with mouse IL12p70 ELISA Kits. The purified cytokine solutions were aliquoted and stored at −20°C. The purified scIL12IL2GMCSF protein was sampled and diluted to 50 ng/ul, then subjected to SDS page electrophoresis under reduced condition.

### Measurement of fusion cytokine bioactivity

Splenocytes from C57BL/6 mice were resuspended in RPMI1640 (2% FBS) with PHA (ebioscience) plus indicated cytokines. IL12, IL2 and GMCSF were purchased from Peprotech. Cells were plated into 96 well plate (6×10^5^/well for experiments in figure 3 and 2×10^5^/well for experiments in figure 5) and cultured for 24 hours. The supernatants were collected and subjected to ELISA measurement of IFNγ using Mouse IFN-gamma Quantikine ELISA Kit (R&D systems).

### Comparison of carboxymethyl cellulose, chitosan and glycerol

10^5^ B16F10 cells were subcutaneously injected into the right flank of mice. Treatment was initiated when the tumor diameter reached around 5mm. 5 ug scIL12IL2GMCSF protein was separately mixed with 50 ul 1% carboxymethyl cellulose, 50 ul 3% chitosan (Chitosan glutamate, Protosan G 213, NovaMatrix) or 60 ul glycerol, generating 100 ul intratumoral injecta. The tumor sizes were daily monitored post intratumoral administration.

### Tumor treatment

10^5^ B16F10, 10^5^ B16F10-rtTA, 2×10^5^ LLC, 10^6^ EL4 or 5×10^5^ MC38 cells were subcutaneously injected into the right flank of C57BL/6 mice. 10^6^ 4T1 or 10^5^ CT26 cells were subcutaneously injected into the right flank of BALB/c mice. Treatment was initiated when the tumor reached 5-9mm in diameter.

For intratumoral administration, cytokine solution was diluted to 40 ul with PBS, and mixed with 60 ul glycerol prior to injection. The glycerol/cytokine solution was slowly injected into tumors using a 29G insulin syringe, avoiding bubbling. In 4T1 tumor treatment, additional injections were conducted as tumor size increased or new nodule appeared.

For intravenous administration, 50 ug recombinant protein was diluted to 200 ul with PBS prior to injection. Five consecutive injections were conducted by once every day (qd), two days (q2d), or three days (q3d). Injection of PBS daily was used as control.

### Tumor sizes and mouse weight were recorded after treatment

### Western blot

2×10^5^ LLC were subcutaneously injected into the right flank of C57BL/6 mice. 200 ug scIL12IL2DiaGMCSF in 200 ul PBS was intravenously injected when tumor grew to 5-8 mm in diameter. Mice were sacrificed at 0, 1 or 2 days post injection. The total protein from tumor tissues were extracted using RIPA. Western blot was carried out following standard procedure, using the antibody rabbit anti-IL12p40 (Abcam).

Immune activation measurement after intravenous administration of recombinant protein 2×10^5^ LLC were subcutaneously injected into the right flank of C57BL/6 mice. 200 ug scIL12IL2DiaGMCSF in 200 ul PBS was intravenously injected when tumor grew to 5-8 mm in diameter. Mice were sacrificed at 0, 1 or 2 days post injection. The IFNγ in serum was detected using Mouse IFN-gamma Quantikine ELISA Kit (R&D systems), and splenocytes were subjected to FACS analysis of immune cell populations.

### Statistics

Statistical analysis was carried out using GraphPad Prism 5 software. Survival curves were analyzed using the log-rank (Mantel-Cox) test. The comparison of growth curves was conducted with two-way ANOVA. p<0.05 was considered to indicate statistical significance, ***p<0.001.

## Acknowledgments

We thank Dr. Wenwen Zeng for experimental advices.

## Funding

There is no external funding supporting this study.

## Author contributions

Jinyu Zhang designed the study, carried out most of the experiments and prepared the manuscript. Xuan Zhao performed experiments including immune cell flow cytometry detection, in vitro activity measurement of the fusion proteins, western blot and serum IFNγ measurement.

## Competing interests

Jinyu Zhang has patents CN201910348357, CN201910351987 and CN201910685007 pending. The other author declares no conflicts of interests.

## Supplementary Materials

**Fig. S1.**
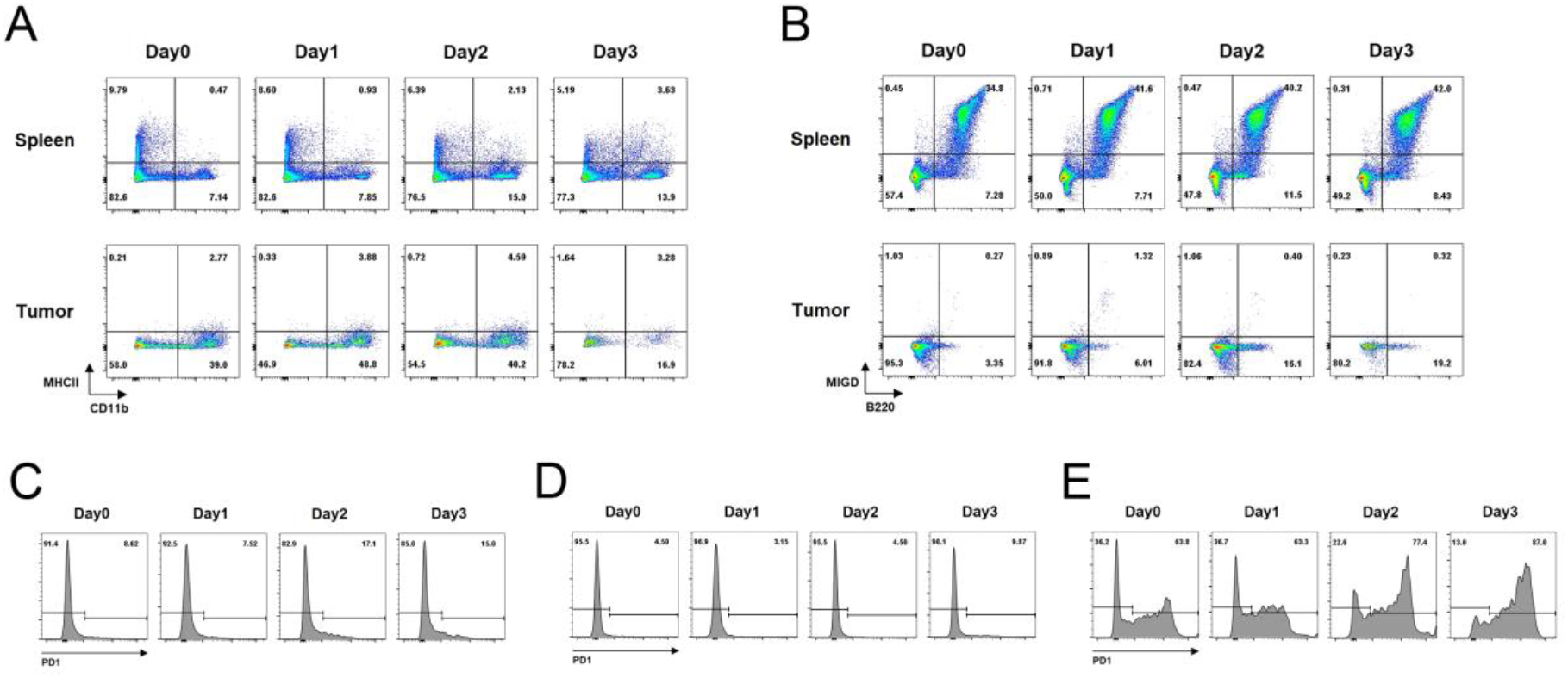
The alterations of lymphocytes during induced expression of dcIL12IL2GMCSF. The inducible B16 cells were subcutaneously inoculated into the flank of C57BL/6 mice. At different times post dox administration, splenocytes and tumor infiltrating lymphocytes were isolated and subjected to flow cytometry analysis. The gated CD45+ cells were analyzed with the groups of CD11b, MHCII (A) or B220, MIGD (B). The PD-1 expression in spleen CD3+/CD4+ (C), CD3+/CD8+ (D) and tumor infiltrating CD3+ cells (E) was detected. The experiments were performed twice with n=2 for each run.

**Fig. S2.**
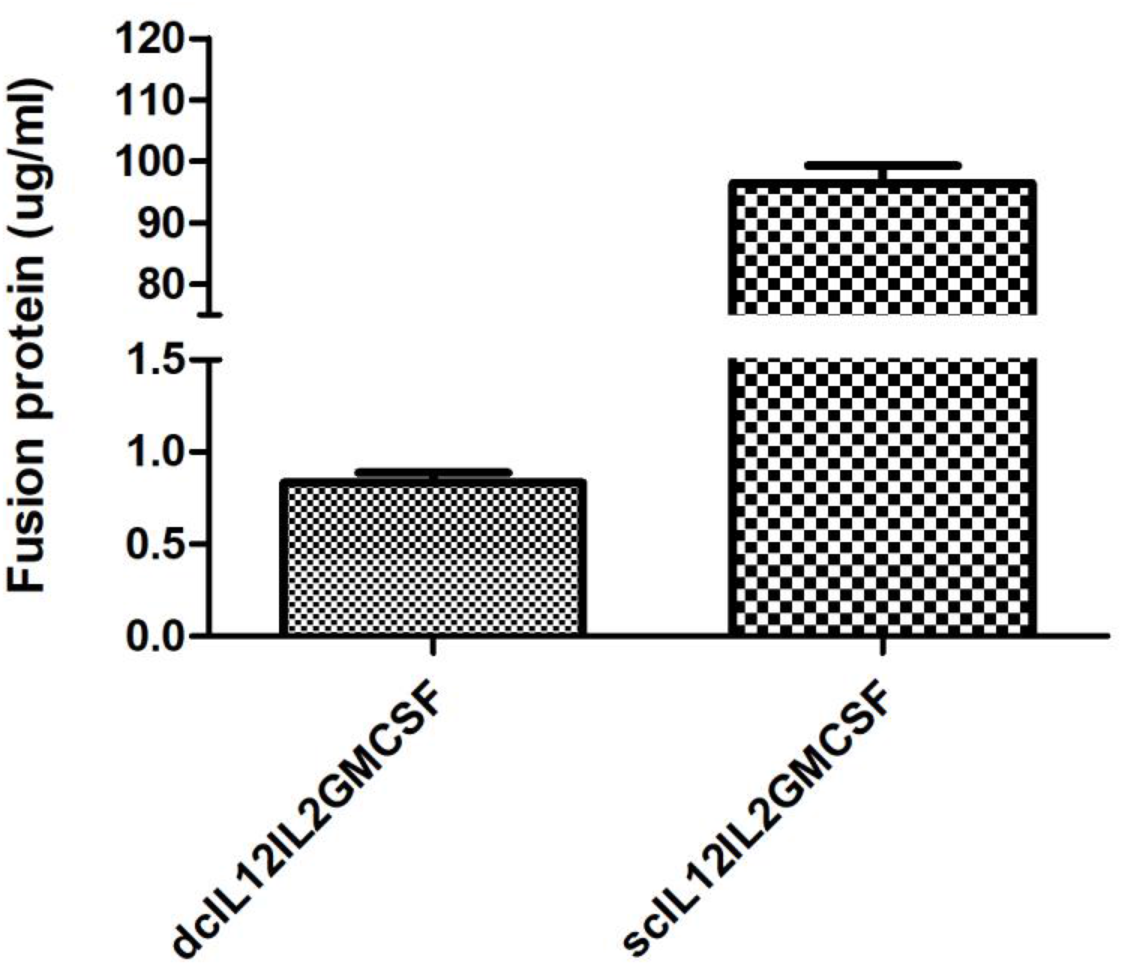
Expression levels of dcIL12IL2GMCSF and scIL12IL2GMCSF. The supernatants of 293(dcIL12IL2GMCSF) or 293(scIL12IL2GMCSF) in CDM4HEK293 culture were collected and subjected to ELISA measurement of recombinant protein level, using mouse IL12p70 ELISA Kits. n=3.

**Fig. S3.**
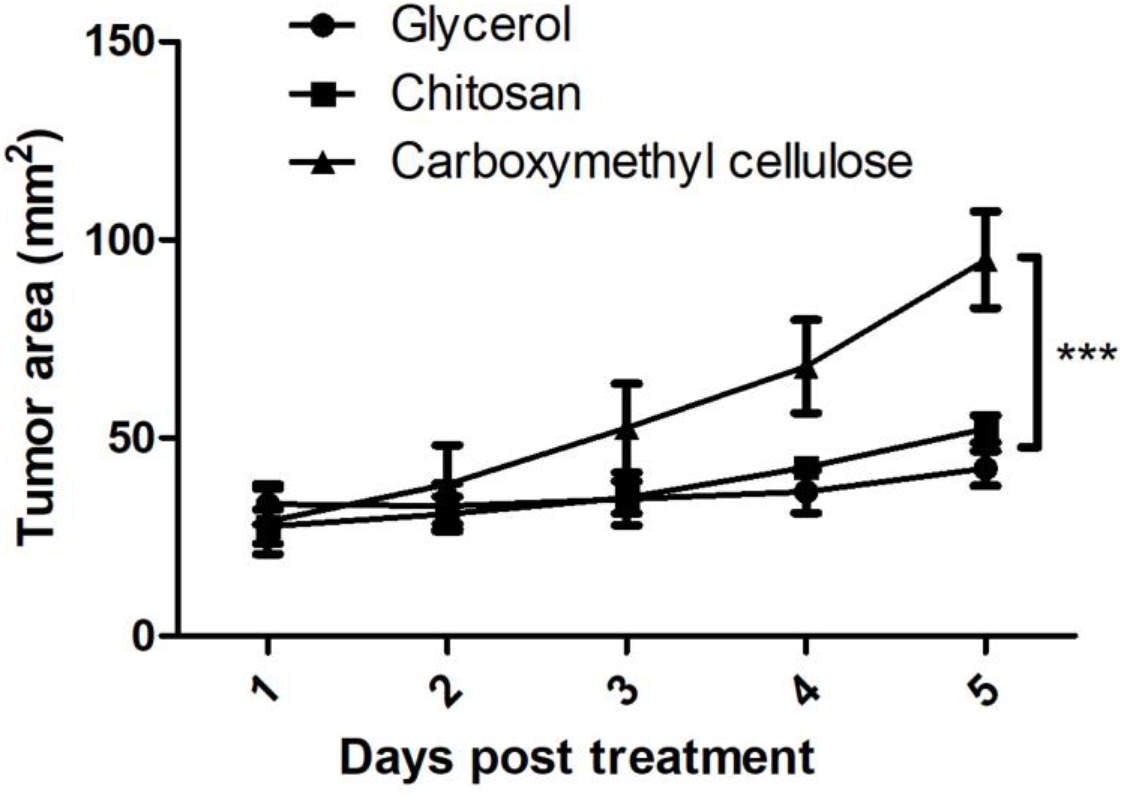
Comparison of therapeutic effects of scIL12IL2GMCSF in different solvents. B16F10 cells were subcutaneously inoculated into the flank of C57BL/6 mice. When tumor reached ~5 mm in diameter, 5 ug scIL12IL2GMCSF in 100 ul 0.5% carboxymethyl cellulose, 1.5% chitosan or 60% glycerol was intratumorally injected. Tumor growth was daily recorded. n=3, ***p<0.001. The experiments were repeated twice.

**Fig. S4.**
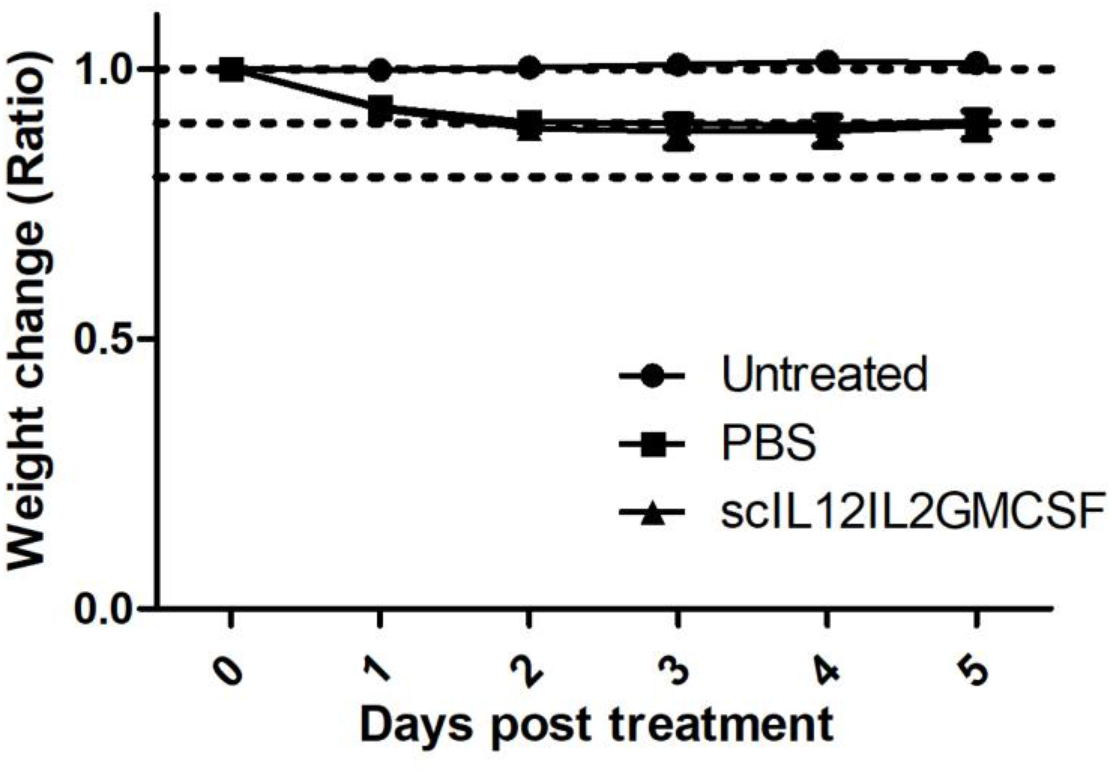
Mouse body weight changes during treatment of B16F10 by intratumoral injection. B16F10 cells were subcutaneously inoculated into the flank of C57BL/6 mice. scIL12IL2GMCSF or PBS in glycerol solution was intratumorally injected into the lesions when tumors reached 5-9 mm in diameter. Body weights were daily recorded and the change percentage was calculated. Untreated tumor bearing mice were used as control. n=5. The experiments were repeated twice.

**Fig. S5.**
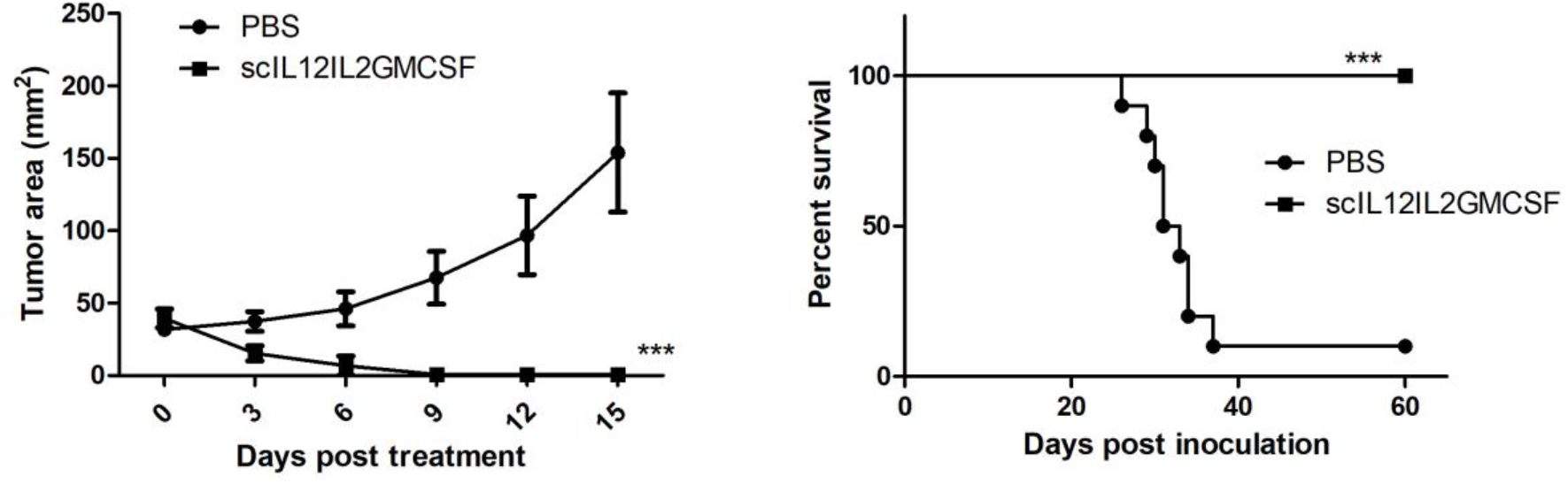
Therapeutic effects of scIL12IL2GMCSF in B16F10-rtTA tumor. B16F10-rtTA cells were subcutaneously inoculated into the flank of C57BL/6 mice. scIL12IL2GMCSF or PBS in glycerol solution was intratumorally injected into the lesions when tumors reached 5-9 mm in diameter. Tumor growth (n=5) and overall survival (n=10) were recorded. ***p<0.001.

**Fig. S6.**
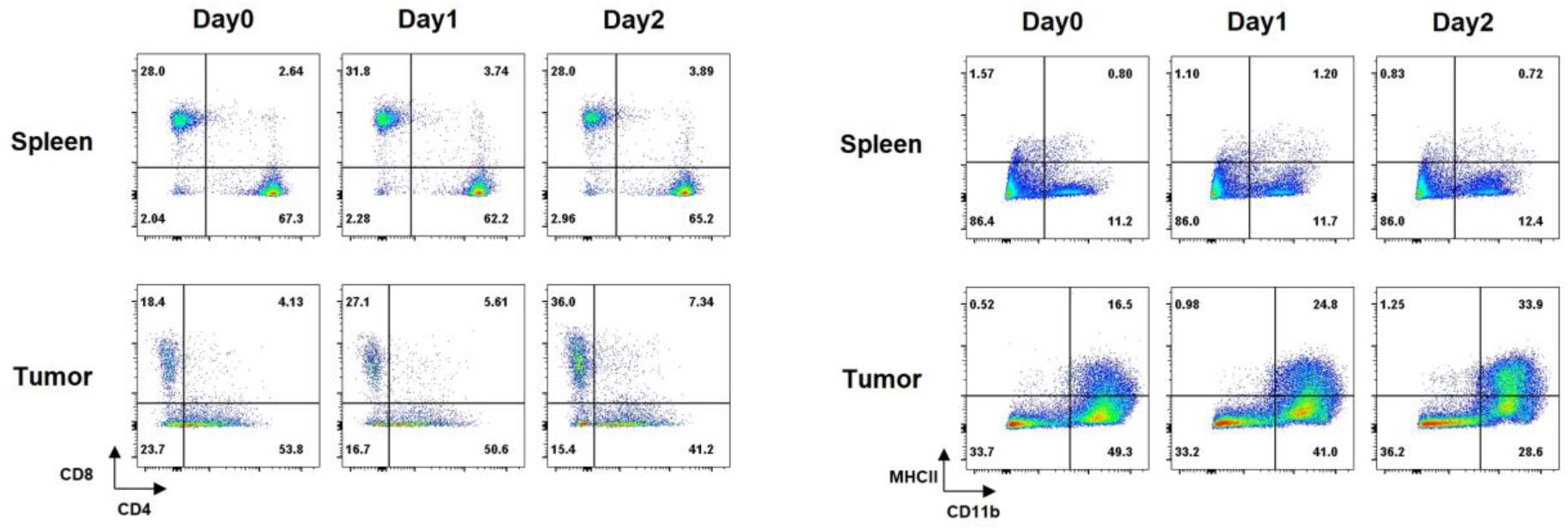
The alterations of lymphocytes after intravenous injection of scIL12IL2DiaNFGMCSF. Mouse LLC tumor cells were subcutaneously inoculated into the flank of C57BL/6 mice. When tumor reached 5-8 mm in diameter, 200 ug scIL12IL2DiaNFGMCSF was intravenously injected into mice. At day 0, 1 or 2 post injection, the splenocytes and tumor infiltrating lymphocytes were subjected to flow cytometry analysis. In gated CD45+ cells, CD11b, MHCII staining group and CD4 and CD8 expression in CD3+ cells were shown. The experiments were performed twice with n=2 for each run.

**Fig. S7.**
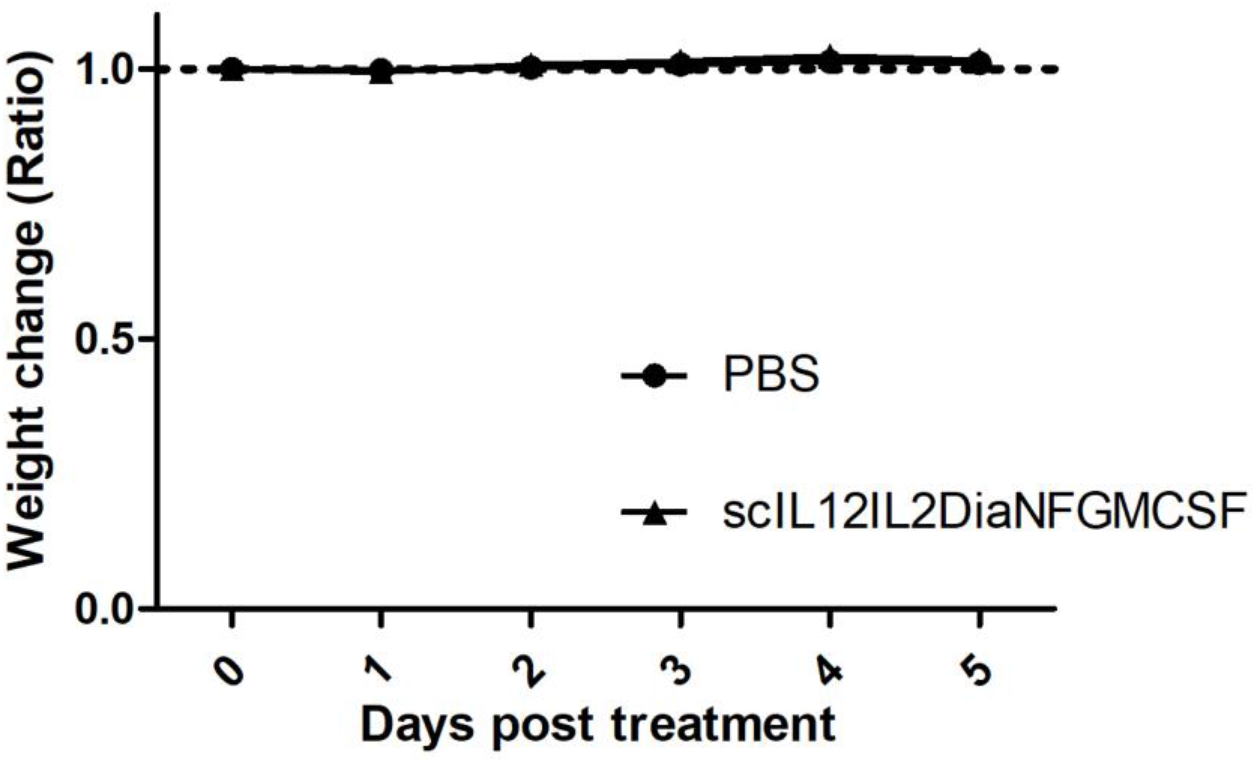
Mouse body weight changes during treatment of B16F10 by intravenous injection. B16F10 cells were subcutaneously inoculated into the flank of C57BL/6 mice. When tumor reached 5-9 mm in diameter, daily intravenous administration of 50 ug scIL12IL2DiaNFGMCSF or PBS was conducted for five days. Body weights were daily recorded and the change percentage was calculated. n=5. The experiments were repeated twice.

**Fig. S8.**
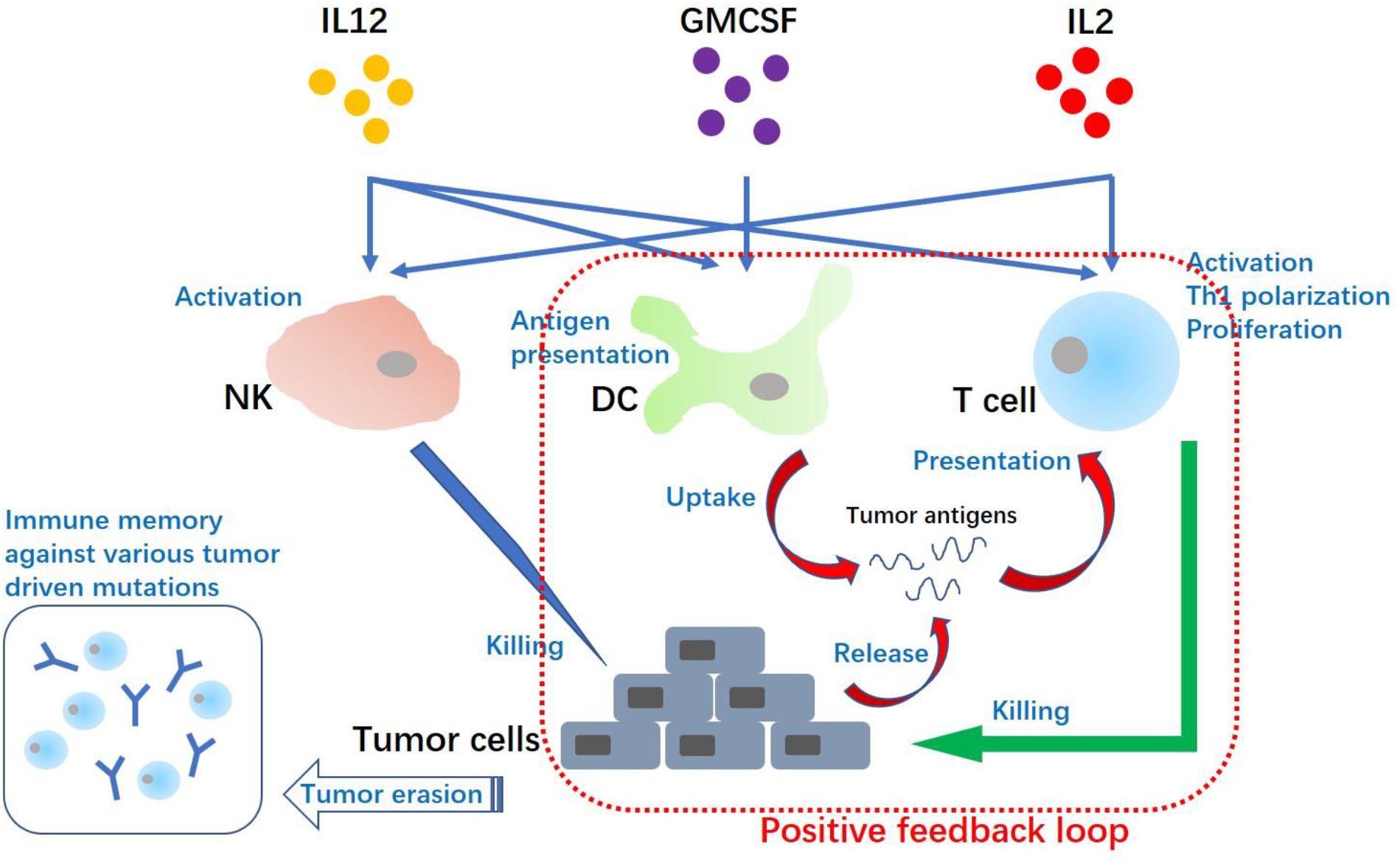
Speculative mechanism of the fusion cytokines in antitumor immunity.

